# Monotreme transcriptomes shed light on evolution of lactation and placentation

**DOI:** 10.64898/2026.08.20.746094

**Authors:** Isabella Wilson, Tahlia Perry, Alexander Stuart, Mish Simpson, Frank Grutzner

## Abstract

Placentation and lactation are hallmark mammalian reproductive traits and key processes facilitating the transfer of nutrients from the mother to the offspring. The basal mammalian lineage of egg-laying monotremes features a short gestation period supported by a simple placenta and an elongated lactation, similar to marsupials. Eutherian mammals by contrast evolved longer gestation facilitated by extensive placentation and shorter lactation. To understand the gene expression underpinning maternal-foetal nutrient transfer across mammals, we generated transcriptome datasets for the monotreme reproductive tract and mammary gland and compared this to expression in representative species from the other mammalian clades: marsupials (Tammar wallaby) and eutherians (mouse). This revealed extensive overlapping gene expression between the therian placenta and monotreme reproductive tract, including the first observation of retroviral gene expression in the echidna reproductive tract. Interestingly, we discovered that transport and development functions supporting foetal growth are expressed in echidna mammary gland and eutherian placenta, likely a signature of the shift from prolonged lactation to longer placentation. Together, this demonstrates that diverse modes of maternal-foetal nutrient transfer are underpinned by similar gene networks across mammals.

## Introduction

Since the emergence and subsequent radiation of mammals, two different maternal-offspring nutrient transfer strategies have evolved to replace the role of egg yolk in supporting early development of offspring^1^. Marsupials and monotremes bear altricial young which are nursed by a lengthy and compositionally complex lactation. Eutherian mammals by contrast mostly evolved longer gestation and shorter lactation periods, facilitated by their complex placentation^2,3^. Interestingly, there is evidence of convergent gene expression between eutherian placentation and marsupial lactation^4^.

The monotremes (platypus and echidna) are the most basal extant mammalian lineage and thus provide unique insight into the evolution of these hallmark mammalian traits. As the only living oviparous mammals, their young are primarily nourished through lactation. Monotremes lack nipples and instead secrete milk from a patch where ducts open onto the flat areola. The mammary glands undergo significant morphological changes during breeding season and lactation^5^, though little is known about this process on a molecular level. Despite the lack of information on the mammary gland, several studies have investigated the composition of monotreme milk. Like marsupials, monotreme lactation is extended (lasting from 114-145 days in the platypus and 160-210 days in the echidna) and may be modulated over this period according to the needs of the young^6^. The milk composition is similar to that of eutherians in a number of ways, being made up of caseins (including α, β, and κ caseins) and whey proteins (including β-lactoglobulin and α-lactalbumin)^7,8^. However, in contrast to eutherian milk, which is high in lactose, monotreme milk predominantly contains oligosaccharides^9^. Echidna milk cell transcriptomes have revealed the expression of monotreme-specific antimicrobial milk proteins, *echAMP* and *MLP*^10,11^, suggesting that monotreme milk serves a protective role as well as providing nutrition. The antimicrobial properties of milk patch secretions are also demonstrated by changes to the resident microbiota during lactation^12^, similar to what occurs in marsupials^13,14^.

While lactation and egg yolk are the main contributions to the development of the offspring, there is nutrient transfer by the reproductive tract during the brief intrauterine gestation period. Monotremes have two separate uteri, only one of which carries an egg during gestation, and in the platypus, only the left side of the reproductive tract can become active^15^. During gestation, monotremes develop an allantoic vitelline placenta from vitellocytes, which are homologous to eutherian trophoblasts^16^. Monotreme placentae do not make contact with the uterine epithelium and are thus non-invasive. However, reproductively active individuals exhibit extensive uterine gland formation, which produce a nutritive secretion that passes through the thin shell coat and into the egg^17–19^. These secretions are thought to be taken up by the placenta, which expands to maximise its surface area (and therefore nutrient exchange potential)^20^. The molecular underpinnings of monotreme placentation are not well understood. Monotremes possess some genes important for placentation, such as *POU5F1*, indicating some capacity for commitment to a trophoblast lineage^21^. Additionally, a number of putative syncytin genes were recently discovered within the echidna genome, of which at least one (*Env-Tac1)* has cell-cell fusion activity^22^. *GCM1*, a transcription factor responsible for regulating syncytin expression in the human placenta, and a master regulator of placental cell differentiation more broadly, is expressed in the active reproductive tract of both the platypus and echidna^23^. Other essential placentation genes such as *PEG10* are absent^24^, potentially explaining the relatively simple nature of the monotreme placenta. However, the expression of other known mammalian placental genes in the monotreme reproductive tract remains unexplored.

Given their shared role in supplying nutrients to the developing offspring, we hypothesised that gene expression in the monotreme reproductive tract and mammary gland may show similarities to the eutherian placenta, despite the evolutionary differences between these organs and species. To investigate this, we generated transcriptome datasets from platypus and echidna reproductive tract, as well as the echidna mammary gland, and compared gene expression to published data in a marsupial (Tammar wallaby) and a eutherian (mouse). This analysis revealed that the monotreme reproductive tract and mammary gland utilise some of the same gene networks as the eutherian placenta to support development of the offspring.

## Materials and Methods

### Overview of datasets

Several transcriptome datasets – both newly generated and from existing studies – were used in this study to compare the gene expression of the reproductive tract, placenta, and mammary gland across mammalian clades. An overview of the datasets can be found in Table 1. For detailed information including sample accessions and dataset validation, see Supplementary File 1.

**Table 1:** Summary of transcriptome datasets used in the present study.

| Species | Tissue | Replicates | Source |
| --- | --- | --- | --- |
| Platypus | Active reproductive tract | 1 | Present study |
|  | Inactive reproductive tract | 1 | Present study |
|  | Liver | 1 | Brawand et al<sup>25</sup> |
|  | Testis | 1 | Brawand et al<sup>25</sup> |
| Echidna | Active reproductive tract | 1 | Present study |
|  | Inactive reproductive tract | 1 | Present study |
|  | Lactating mammary gland | 2 | Present study |
|  | Non-lactating mammary gland | 1 | Present study |
|  | Liver | 1 | Zhou et al<sup>26</sup> |
|  | Testis | 1 | Zhou et al<sup>26</sup> |
| Wallaby | Gravid endometrium | 2 | Dudley et al<sup>27</sup> |
|  | Non-gravid endometrium | 2 | Dudley et al<sup>27</sup> |
|  | Liver | 1 | Cortez et al<sup>28</sup> |
|  | Testis | 1 | Cortez et al<sup>28</sup> |
| Mouse | Gravid endometrium | 1 | Siewiera et al<sup>29</sup> |
|  | Placenta | 1 | ENCODE |
|  | Lactating mammary gland | 4 | Fu et al<sup>30</sup> |
|  | Non-lactating mammary gland | 2 | Fu et al<sup>30</sup> |
|  | Liver | 1 | ENCODE |
|  | Testis | 1 | ENCODE |

### Sample collection

Platypus and echidna reproductive tract and echidna lactating mammary gland samples were collected and snap-frozen in liquid nitrogen before being stored at - 190°C (AEEC permit number R.CG.07.03, AEC permit numbers S-49-2006 and S-032-2008). Non-lactating echidna mammary gland sample was collected by Southern Koala and Echidna Rescue Ltd (permit numbers: SH2545906E and RC2106738V) and stored in RNA*later* (Invitrogen).

### Library preparation and sequencing

Tissue samples were homogenised by grinding in a mortar and pestle with liquid nitrogen. Total RNA was extracted using the RNeasy Plus Micro Kit (Qiagen) according to the manufacturer’s instructions. Extracted RNA was sent to the Australian Genome Research Facility (AGRF) where quality was assessed using the Agilent TapeStation 4200 and quantity was determined using the Promega QuantiFluor RNA System. mRNA was isolated using PolyA+ enrichment (Illumina Stranded mRNA Prep kit). After further quality and quantity checks, libraries were normalized and pooled in an equimolar fashion before being sequenced on an Illumina NovaSeq X Plus. Sequencing data will be made available on BioProjects upon publication.

### Data processing

Trimming and quality control was performed using TrimGalore^31^. Trimmed reads from each species were aligned to their respective genomes - platypus (mOrnAna1.pri.v4)^26^, echidna (mTacAcu1.pri)^32^, wallaby (mMacEug1.pri_v2)^33^, and mouse (mm10)^34^ - using STAR (V. 2.7.10b_alpha_230301)^35^. Pre-mapped mouse transcriptome data was obtained from ENCODE for placenta (ENCSR000BZP), liver (ENCSR216KLZ), and testis (ENCSR266ESZ). Gene counts were generated using the featureCounts function in the Rsubread R package (v. 2.20.0)^36,37^.

### Comparative transcriptomics

To make datasets from different species comparable, we restricted our analysis to 1:1 orthologs. These were identified using Orthofinder (v. 3.1.0)^38^ using the longest isoform of each gene for each species. Data analysis was carried out using the EdgeR R package (v. 4.4.1)^39^; full data analysis code is available at https://github.com/isa-wilson/monotreme-transcriptome. Where necessary, mouse Ensembl gene IDs were converted to gene symbols using the biomaRt R package (v. 2.62.0)^40^. MDS plotting based on CPM was used to visualise sample similarity and identify potential outlier datasets; due to high sample similarity, active and inactive reproductive tract and endometrium samples were merged (further detail regarding data validation is available in Supplementary File 1). To assess similarity of gene expression between each transcriptome dataset, we performed a series of pairwise correlations (Spearman’s rank correlation coefficient). First, TPM was calculated for each sample, after which biological replicates were merged by mean TPM. The samples were ranked by TPM, and pairwise Spearman correlations were calculated using the R stats cor() function.

Spearman correlations were visualised using a heatmap ggplot2 (v. 3.5.2)^41^. Differential expression analysis was performed on echidna and mouse mammary gland tissues using the EdgeR quasi-likelihood pipeline. Genes were considered significantly differentially expressed when q ≤ 0.05. Of the significantly differentially expressed genes, those with a log-fold-change > 1 were considered as showing higher expression, and those with a log-fold-change < -1 were considered as showing lower expression.

Full results for differential expression analysis can be found in Supplementary File 2. For gene ontology analysis, genes were designated as “expressed” using a threshold of CPM > 1. Lists of expressed genes are available in Supplementary File 3. InteractiVenn (https://www.interactivenn.net)^42^ was used to visualise and compare overlapping sets of expressed genes between datasets. Gene set enrichment analysis was performed using the GO Biological Process and Reactome Pathway databases via ShinyGO v. 0.85.1 (https://bioinformatics.sdstate.edu/go/)^43^ and the Reactome Analyse Gene List online tool v. 95 (https://reactome.org/PathwayBrowser/#TOOL=AT)^44^. Detailed statistics for gene set enrichment analysis is available in Supplementary File 4.

### Syncytin expression analysis

Analysis of syncytin gene expression in echidna tissues was performed using Rsubread featureCounts. A custom annotation file was created based on the env ORF gene coordinates provided in Kitao et al^22^. The EdgeR was used to calculate CPM values and ggplot2 was used for data visualisation.

## Results

### Shared gene expression suggests functional similarities between the monotreme reproductive tract and the therian placenta

The monotreme reproductive tract supplies nutrients to developing embryos, resembling the function of the therian placenta. We tested for similarities in the overall gene expression of these tissues and compared this to the gene expression of other organs including reproductive tract, liver and testis in therian mammals. We conducted a series of Spearman correlation tests based on the normalised expression (TPM) of 1:1 orthologs between the echidna, platypus, mouse, and wallaby (Figure 1A). When comparing the echidna reproductive tract to the mouse tissues, the highest level of similarity was with the mouse placenta (ρ = 0.64) – marginally higher than with the equivalent tissue (mouse reproductive tract, ρ = 0.63). Platypus reproductive tract also exhibited similarity to the mouse placenta (ρ = 0.62), though less than to mouse reproductive tract (ρ = 0.65). Surprisingly, the wallaby placenta was equally as similar to the mouse placenta (ρ = 0.64), though this lower-than-expected similarity may have been due to a low read assignment rate for the wallaby placenta datasets (Supplementary Table S1). Together, this demonstrated shared gene expression between the monotreme reproductive tract and eutherian placenta.

**Figure 1:**
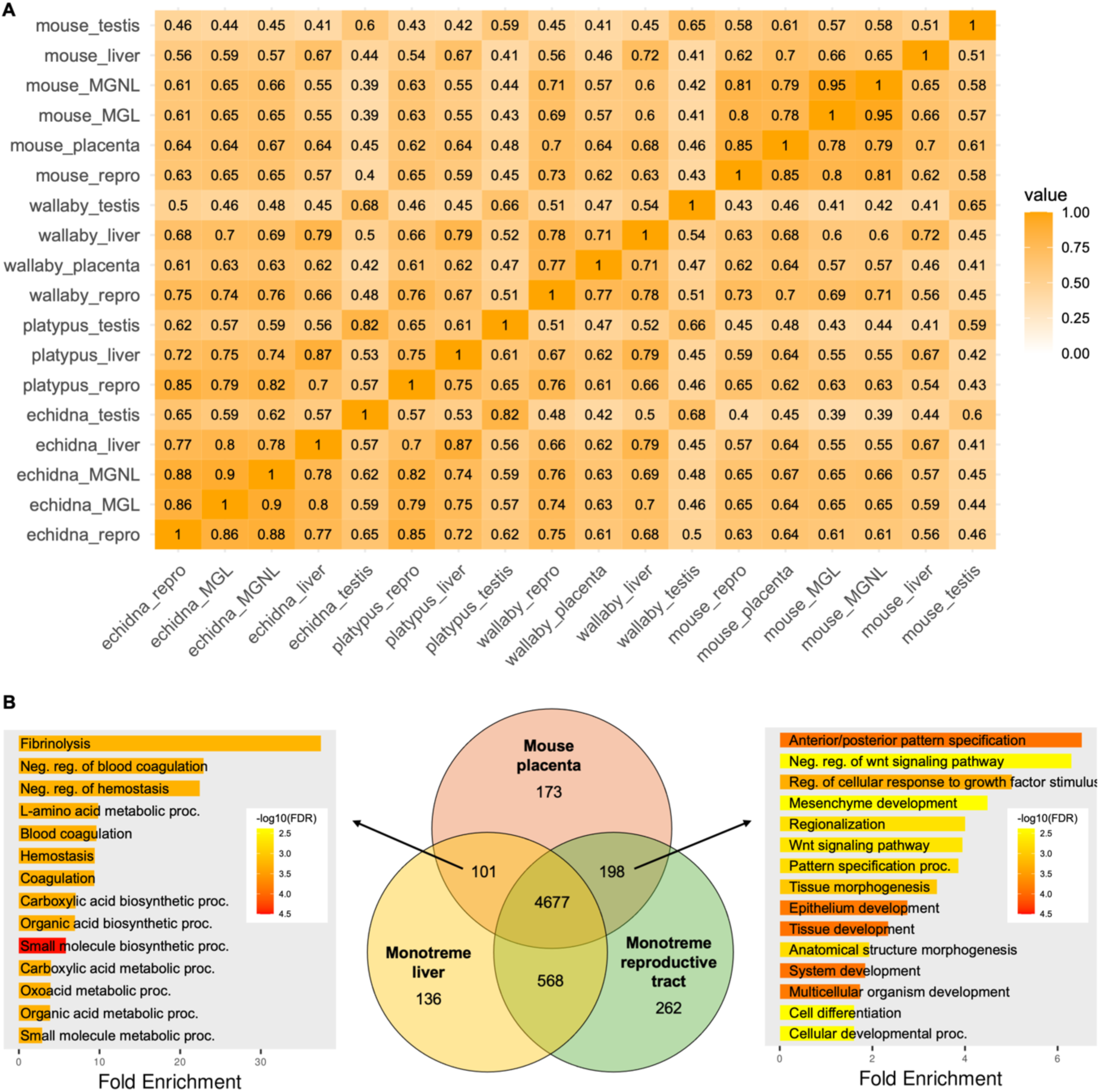
Gene expression in monotreme reproductive tract and therian placenta. (A) Heatmap comparing similarities in gene expression between tissues from platypus, echidna, wallaby, and mouse. Numbers indicate Spearman correlation coefficients; dark orange indicates a high degree of similarity. Abbreviations: repro = reproductive tract, MGL = lactating mammary gland, MGNL = non-lactating mammary gland. (B) Functional analysis of overlapping gene expression between the mouse placenta, monotreme reproductive tract, and monotreme liver. Bar plots show enrichment analysis using the GO Biological Process database; the top 15 most significant terms (ordered by fold enrichment) are displayed.

This shared expression may reflect general cellular functions rather than placenta-specific functions, especially given that the similarity between the mouse placenta and monotreme liver samples was relatively high (ρ = 0.64). To gain functional insight into gene expression shared between these tissues, we performed gene set enrichment analysis (Figure 1B). We observed highly divergent functional profiles when comparing shared expression in the mouse placenta and monotreme reproductive tract (total 198 genes) to shared expression in the mouse placenta and monotreme liver (total 101 genes) (Figure 1B). Genes shared by the mouse placenta and monotreme liver were enriched for liver-associated metabolic processes including the urea cycle and hemostasis. In contrast, genes shared by the mouse placenta and monotreme reproductive tract genes were enriched for developmental processes. The mouse placenta-monotreme reproductive tract ontologies with the highest fold-enrichment pertained to early embryogenesis, such as anterior/posterior pattern specification (q = 0.00012), mesenchyme development (q = 0.0041), and branching morphogenesis (q = 0.0041). There was also enrichment for cellular responses to growth factor (q = 0.0042) and the WNT signalling pathway (q = 0.0021). Pathway analysis (Reactome) revealed that the developmental ontologies pertained to neurogenesis, nephrogenesis, ectoderm formation, and the epithelial-mesenchymal transition, with these processes being underpinned by HOX and WNT signalling pathways as well as *TFAP2A*. A similar pattern was observed when comparing the monotreme reproductive tract to the wallaby placenta (Supplementary Figure S3.) The shared gene expression between the monotreme reproductive tract and therian placenta, particularly in pathways relating to development and placentation, provides evidence of overlapping roles for these tissues in supporting the development of the embryo.

Syncytins are genes of retroviral origin that have been captured for placenta function in eutherian and marsupial lineages^45,46^. Given the broad transcriptional similarity between the monotreme reproductive tract and therian placenta, we investigated expression of retroviral genes in the echidna reproductive tract. We found that 24 of the 121 putative syncytin ORFs within the echidna genome were expressed in the echidna reproductive tract (Figure 2). In particular, *Env-Tac1* was very highly expressed (CPM = 120) in the echidna reproductive tract and was co-expressed with its receptors *SLC1A4* and *SLC1A5*. Another ORF, *Env-Tac4.1*, exhibited high levels of reproductive tract expression (CPM = 58) relative to other tissues examined. Several syncytin ORFs were also expressed in the liver and testis at similar levels to what has previously been reported^22^.

**Figure 2:**
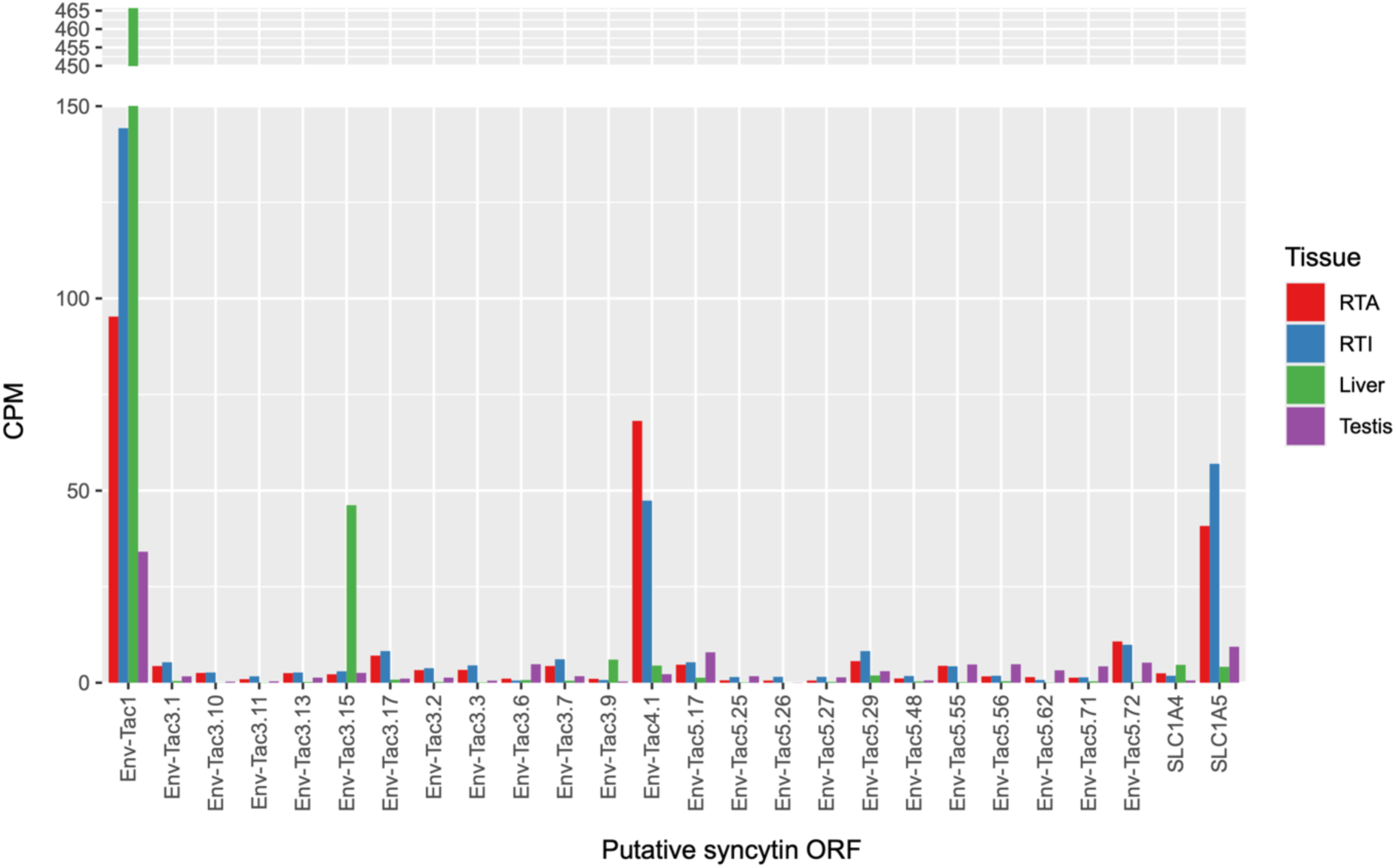
Expression of putative syncytin open reading frames (ORFs) in echidna tissues. Expression was calculated using multimapping reads and normalised using counts per million (CPM).

### Molecular links between monotreme lactation and eutherian placentation

In monotremes and marsupials, the young hatch or are born at an early developmental stage relative to eutherian mammals. Lactation therefore performs a similar role to the placenta in eutherian mammals: supplying nutrients to support early development. Previous work has provided evidence that Tammar wallaby mammary gland and mouse placenta recruited the same gene networks to support convergent functions of these organs^4^. We found that the gene expression of the echidna mammary gland was almost as similar to the mouse placenta (ρ = 0.64) as it was to mouse mammary gland (ρ = 0.65) (Figure 1A). To gain an understanding of the lactation-associated gene expression of the echidna mammary gland, we performed gene-set enrichment analysis, excluding genes also expressed in the liver to focus on mammary gland-specific function (Figure 3A). This revealed enrichment for developmental processes including neurogenesis (q = 1.5E-7) and epithelium development (q = 0.00016). Interestingly, this resembled the gene ontologies that we previously found were shared between the monotreme reproductive tract and mouse placenta - specifically, nervous system development and *TFAP2A* signalling.

**Figure 3:**
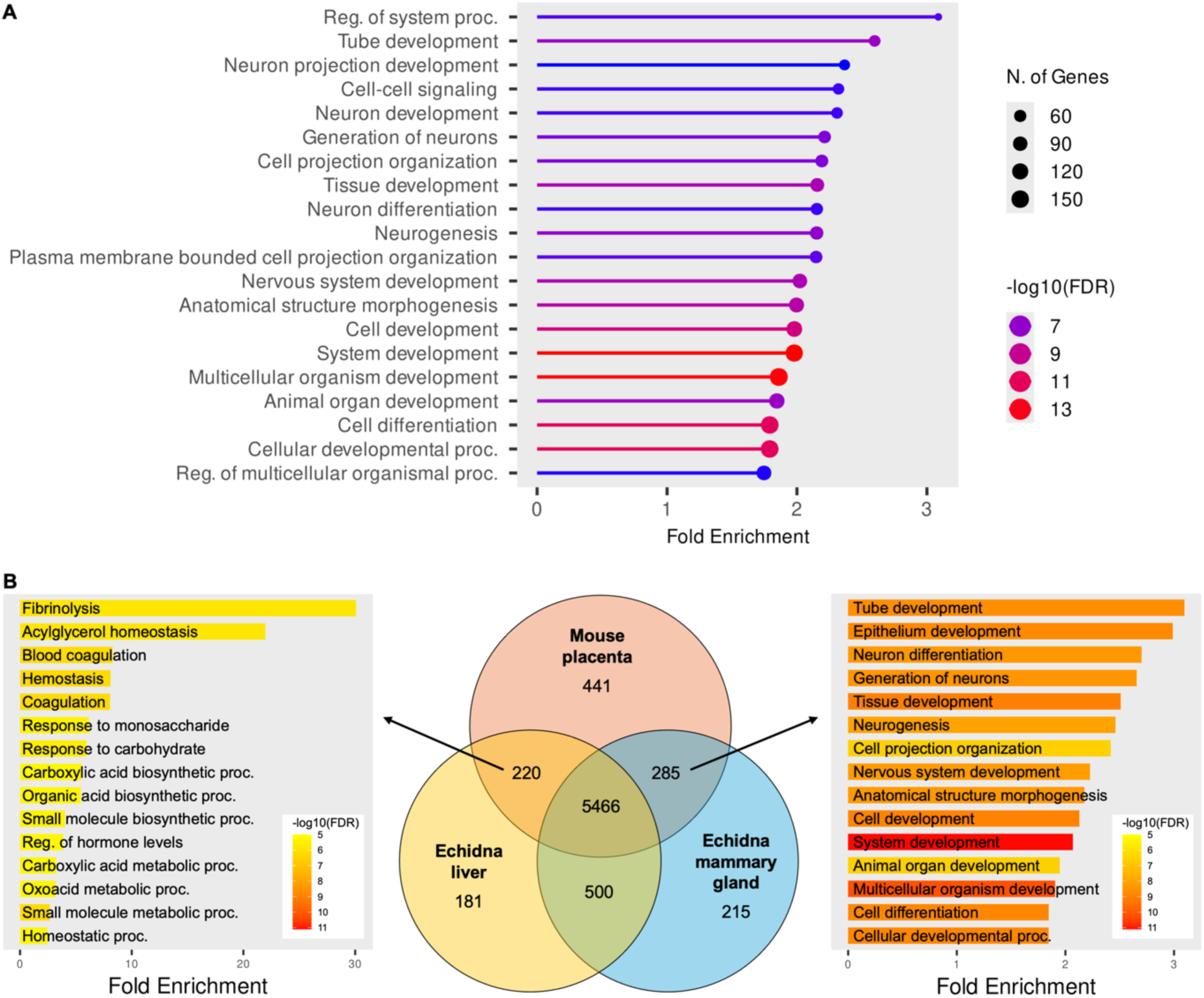
Overlapping functions of the echidna mammary gland and mouse placenta. (A) Ontology of genes expressed in the echidna mammary gland during lactation, excluding liver-expressed genes. (B) Ontological comparison of genes shared by the mouse placenta and monotreme mammary gland with genes shared by the mouse placenta and echidna liver.

To further analyse potential shared expression patterns between the monotreme mammary gland and eutherian placenta, we compared overlapping gene expression between the echidna mammary gland and mouse placenta, also including the echidna liver as a point of comparison (Figure 3B). We found that the ontologies of these gene sets differed greatly. Genes expressed by the mouse placenta and echidna liver were enriched for homeostatic and metabolic processes. By contrast, genes with shared expression in the mouse placenta and echidna mammary gland were enriched for developmental processes, with the highest fold-enrichment ontologies being sensory organ development (GO:0007423; q = 1.8E-6), tube development (GO:0035295; q = 2.3E-9), and epithelium development (GO:0060429; q = 0.0001). There was also enrichment for several ontologies related to neurogenesis as well as vasculature development (GO:0001944; q = 9.77E-6). There was also significant enrichment for placenta development (GO:0001890; q = 0.012) and we identified several placentation genes expressed in the echidna mammary gland and mouse placenta (*STC1*, *PTGS2*, *GJB2*, *GJA1*, *BMP7*, *CDKN1C*, *VASH1*, *GRHL2*, *OVOL2*). To investigate the shared use of pathways supporting foetal development across mammalian evolution, we analysed genes that were expressed in the lactating echidna mammary gland and mouse placenta, but not in the lactating mouse mammary gland (Supplementary Figure S4). This set of 240 genes was significantly enriched for transport functions including import (GO:0098657; q = 0.0002), endocytosis (GO:0006897; q = 0.001) and secretion (GO:0032940; q = 0.002). Additionally, there was enrichment for developmental pathways including neurogenesis (GO:0022008; q = 0.002) and multicellular organism development (GO:0007275; q = 0.002). Overall, our findings show co-option of similar gene networks supporting foetal development in the echidna mammary gland and eutherian placenta.

### Divergence of echidna and mouse mammary gland gene expression

In light of the differences in the maternal-offspring nutrient transfer strategies between monotremes and eutherians, we investigated gene expression underpinning lactation in these species. We performed differential expression analysis between the mouse and echidna mammary gland, identifying 1408 genes showing higher expression in the echidna relative to the mouse and 1353 genes showing lower expression in the echidna relative to the mouse (Figure 4A). The genes showing higher expression in the echidna were significantly enriched only in mitochondrial translation (q = 0.020). In contrast, genes showing lower expression in the echidna relative to the mouse were significantly enriched for over 180 ontologies, of which the highest fold-enrichment terms related to transcription and RNA metabolism (Supplementary Figure S6). Pathway analysis (Reactome) revealed enrichment for processes related to the immune system (Figure 4B). In particular, innate immunity ontologies showed lower expression in the echidna, including activation of toll-like receptors and interleukin-17 signalling. This gene set was also enriched for MAPK activation (q = 3.6E-05) and cellular senescence (q = 3.5E-05), two processes important for mammary gland development and involution. Overall, the pathways with higher expression in the echidna reflected general cell function, whereas those with lower expression were associated with lactation.

**Figure 4:**
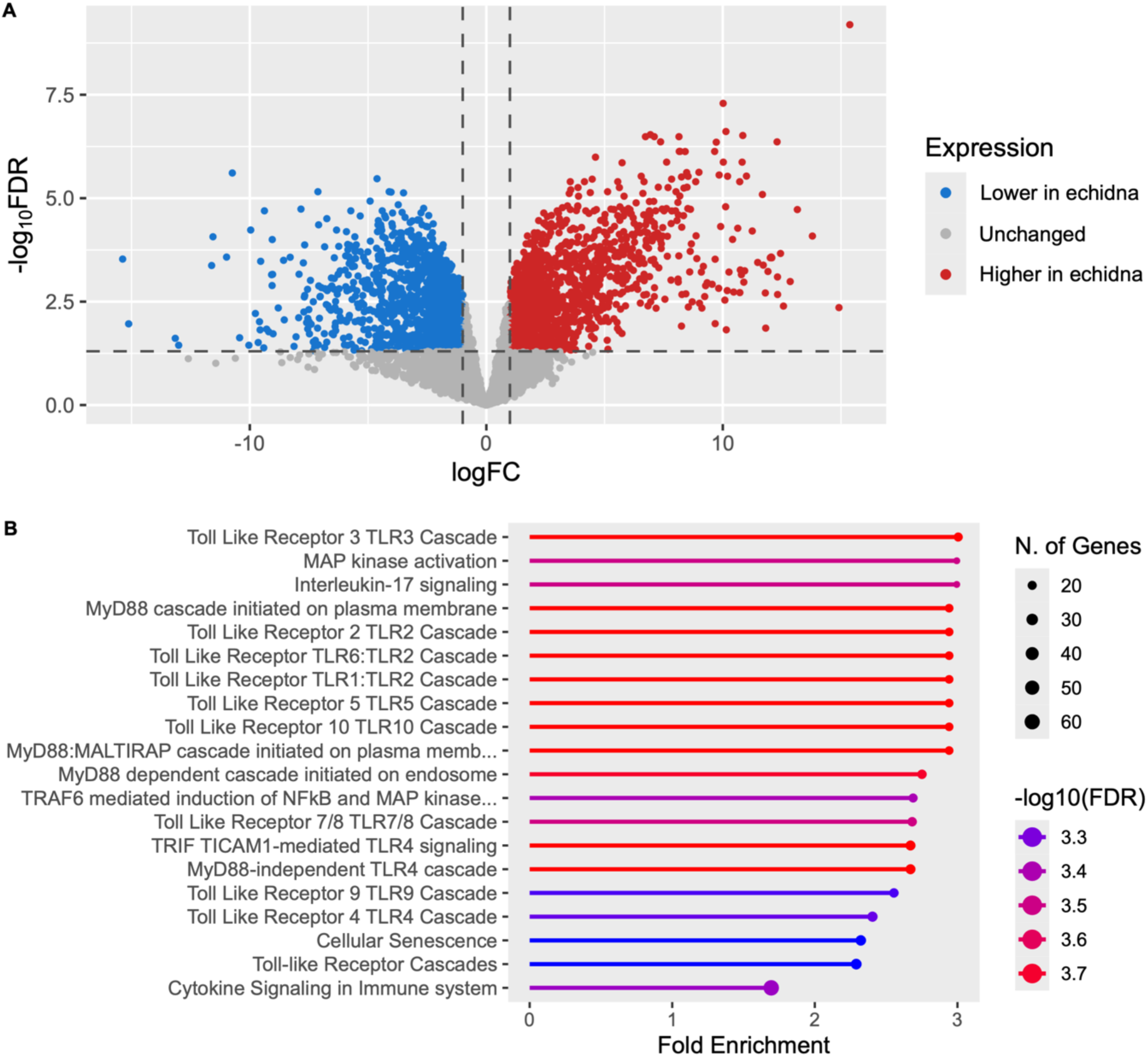
Differential expression analysis of the echidna and mouse mammary gland. (A) Volcano plot showing relative expression of genes in the echidna and mouse. Gene set enrichment analysis of genes with lower expression in the echidna using the Reactome database. Top 20 pathways are displayed in order of fold enrichment.

## Discussion

The unique reproductive biology of monotremes, together with their deep divergence from other mammals, provide important insight into the evolution of lactation and placentation in mammals. Monotreme reproduction features a short intrauterine gestation supported by a simple placenta. Hatching occurs at a very early stage of development followed by a long lactation period. Here, we generated novel transcriptomic data from the platypus and echidna reproductive tract, as well as the echidna mammary gland during lactation. Our analysis revealed expression of similar gene networks between these monotreme tissues and the eutherian placenta.

Whole-transcriptome comparisons revealed hundreds of genes with shared expression between the mouse and wallaby placentas and the monotreme reproductive tract. These shared genes reflected broad functions in development rather than placentation-specific processes, and were enriched for HOX, WNT, and TFAP2A signalling. These pathways have well-characterised roles in eutherian placenta development, being involved in trophoblast differentiation and proliferation, and placenta vascularisation^47–50^. Similar processes take place in the monotreme reproductive tract to produce the secretion of nutritive fluids during gestation, namely increased vascularisation and the growth of more and larger endometrial glands^17,18^. HOX, WNT, and TFAP2A signalling also function in the adult eutherian uterus in the development of endometrial glands, the proliferation of new endometrial tissue and blood vessels each menstrual cycle, as well as being important for embryo implantation^51–53^. Similar gene expression between these monotreme and eutherian tissues highlights that fundamental pathways underpin maternal support for development across mammals. Future research, particularly single-cell sequencing of the monotreme reproductive tract, could further differentiate expression of these pathways in different layers of the endometrium and in the context of maternal-foetal nutrient transfer during monotreme gestation.

Syncytins are retroviral-derived genes with fusogenic and immunosuppressive functions that have been co-opted into placental function in several eutherian lineages independently^54^. More recently, syncytin genes have been identified and characterised in monotremes^22^. We found that 24 of the 121 known syncytin ORFs within the echidna genome were expressed in the echidna reproductive tract. The most highly expressed of these was *env-Tac1*. The cell-cell fusion activity of this ORF is mediated by receptors SLC1A4 (ASCT1) and SLC1A5 (ASCT2)^22^, which we found were co-expressed with *env-Tac1* in the echidna reproductive tract. In eutherians with invasive placentae, the fusogenicity of syncytin genes plays a critical role by causing trophoblast cells to fuse together, forming the multinucleate syncytiotrophoblast layer that invades the maternal epithelium and mediates nutrient transfer. Retroviral genes are important for placenta evolution more broadly; the co-option of regulatory elements derived from endogenous retroviruses likely played a role in establishing the ancestral trophoblast cell type, and the independent integration of retroviruses in different lineages facilitated the morphological diversity of mammalian placentas^47,55,56^. Previous work using platypus endometrial expression data has shown that transposons, another type of mobile genetic element, were important for the recruitment of existing genes into endometrial expression in the mammalian ancestor and to a larger extent in the eutherian ancestor, contributing to the evolution of uterine decidualisation and pregnancy in mammals^57^. Echidnas lack an invasive placenta, so the presence and expression of syncytins suggests a different function in the echidna reproductive tract. Co-expression of *env-Tac1* with its entry receptors *SLC1A4* and *SLC1A5* is indicative of cell-cell fusion activity, though histological studies of echidna uterine tissue do not show a syncytium^17^. Reports of retroviral *env* gene expression in the reproductive tracts of other species are limited. In humans, *syncytin-1* is typically silenced in the endometrium, with examples of pathological endometrial expression in patients with endometriosis or endometrial cancer^58,59^. Research in eutherian mammals has shown that apart from being fusogenic, retroviral *env* genes play a range of other roles including immunosuppression, protection against exogenous viral infections, and inhibition of apoptosis^60,61^. Future studies should investigate which of these functions are performed by the retroviral *env* genes expressed in the echidna reproductive tract.

Similarities in gene expression and function have previously been described between the eutherian placenta and the Tammar wallaby mammary gland. These two tissues share genes enriched for nutrient transfer, immunity, and embryonic development, suggesting that lactation in marsupials parallels the function of the eutherian placenta in supporting late foetal development^4^. Our study demonstrates that echidna mammary gland gene expression also shows a high degree of similarity to the mouse placenta. Like with the Tammar wallaby, the genes shared between the echidna mammary gland and the mouse placenta have functions in embryonic development, specifically in epithelium development and neurogenesis. Epithelium development is a critical process in both the placenta and the mammary gland; during lactation, the mammary epithelium undergoes significant remodelling to form the milk-producing alveoli, whereas during placentation, the epithelial trophoblast layer grows and differentiates into the various cell types that invade the maternal decidua^62,63^. It was unexpected to find enrichment for neurogenesis in genes shared by the echidna mammary gland and mouse placenta as the placenta lacks nerves^64^. It is possible that these genes were involved in neuroendocrine signalling rather than neurogenesis. In rodents, placental neuroendocrine signalling can act locally to promote placental growth; in the foetus where it impacts foetal brain development; or in the maternal brain where it has been linked to the priming of maternal caregiving behaviours^65^. Alternatively, the shared neurogenesis pathways that we observed may reflect the production of neurogenic factors for transport into the developing foetus, rather than supporting local neurogenesis within the mammary gland and placenta^66,67^. When specifically comparing gene expression supporting foetal development in each species (i.e. excluding genes expressed in the mouse mammary gland), we found shared transport and development processes, similar to what has been previously observed between the Tammar wallaby mammary gland and the eutherian placenta^4^. Together, this suggests that the use of these gene networks shifted from the mammary gland to the placenta after the emergence of therian mammals, as placentation replaced lactation as the dominant mode supporting foetal development.

Lactation has changed dramatically over the course of mammalian evolution. Monotremes are thought to most closely resemble the lactation of early mammals^68^, lacking nipples and instead secreting milk from the milk patch. Like marsupials, their milk is compositionally complex and changes throughout their long lactation to support the developmental needs of their offspring. Eutherians, by contrast, typically have a shorter lactation period because the placenta has already provided substantial nutritive support by the time of parturition which occurs at a much later stage of development. To gain insight into the pathways involved in these contrasting lactation strategies, we performed the first differential expression analysis between the mammary gland of a monotreme (the echidna) and a eutherian (mouse). This revealed that genes upregulated in the mouse were enriched for lactation-associated ontologies and pathways, specifically MAPK activation, cellular senescence, and innate immunity. The upregulation of innate immunity pathways in the mouse mammary gland was unexpected given the importance of milk in protecting altricial echidna puggles that lack a functional immune system^69^. One limitation of comparative transcriptomics between different species is that the genes examined must be restricted to 1:1 orthologs to allow comparison. As such, lineage-specific genes - such as those that encode the monotreme-specific antibacterial milk proteins echAMP and MLP – were excluded from our analysis. It is possible that the immune function of lactation in echidnas is shaped by these monotreme-specific genes, rather than the TLRs that we found were upregulated in the mouse. Alternatively, the reduced expression of innate immune genes in the echidna mammary gland may reflect different stages of lactation; the mouse samples were collected very early in lactation (day 2) whereas the echidna samples were likely collected mid-lactation. Future research with sampling at defined timepoints could provide deeper insight into the divergent lactation strategies of monotremes and eutherians. Additionally, current work establishing primary mammary cultures from the echidna (manuscript in preparation)^70^ will provide a powerful model system allowing specific insights into the formation of the mammary gland as well as the evolution of mammary gland expression across mammals.

## Conclusion

The unique reproductive biology and evolutionary position of monotremes make them key species in which to study gene expression during lactation and placentation. To better understand the molecular underpinnings of maternal-offspring nutrient transfer in the monotremes, we generated transcriptome datasets from several monotreme tissues - including the first available echidna reproductive tract transcriptome - and compared gene expression to a marsupial (Tammar wallaby) and a eutherian (mouse). We identified genetic similarities between the eutherian placenta and monotreme reproductive tract, including the expression of putative syncytins, adding to the evidence that the monotreme reproductive tract supports the embryo during the short intrauterine gestation. We also found overlapping expression of genes involved in development and transport between the eutherian placenta and monotreme mammary gland, suggesting that monotreme lactation serves a comparable role to the eutherian placenta in supporting early development. Overall, we found that lactation and placentation recruited similar gene networks in different mammals to support development of the offspring. Access to material from different stages of lactation and placentation in future will allow more detailed analysis of the foetal component of the monotreme placenta to further investigate pre-parturition nutrient transfer in these unique species.

## Acknowledgements

The authors acknowledge all members of the Grützner laboratory for their helpful discussions and feedback throughout this project. We acknowledge the staff at Southern Koala and Echidna Rescue Ltd and Dr Allison Crawley for their help with sample collection. I.W. was supported by a University of Adelaide Research Scholarship.

## Data availability

Raw sequencing data will be made available on BioProjects upon publication. Code used for data analysis and visualisation is available on Github: https://github.com/isa-wilson/monotreme-transcriptome.

