## Supplementary file 1 for "Monotreme transcriptomes shed light on evolution of lactation and placentation"

**Dataset overview and validation**

We generated seven novel monotreme transcriptome datasets from platypus and echidna reproductive tract and mammary gland with an average library size of 26,175,699 reads. We sought to compare our data to equivalent transcriptome datasets from a marsupial (Tammar wallaby) and a eutherian (mouse); see Table 1 for an overview of the samples used. Due to low read assignment rates, we excluded the wallaby mammary gland datasets from further analysis. MDS plotting (Figure S1A) showed that the wallaby placenta clustered separately to all other samples; however, given that the read assignment rate was on average above 50%, we elected to retain these samples while interpreting results cautiously.

Given that this study presents the first characterisation of the echidna reproductive tract transcriptome and new platypus data additional to an existing endometrial transcriptome dataset (see Lynch et. al. 2015), we sought to gain a general understanding of the gene expression of the monotreme reproductive tract. MDS plotting showed that the platypus endometrium sample clustered more closely with the echidna samples than the platypus samples; as such, we excluded this sample from further analysis. Additionally, the samples clustered according to the individual from which they were derived rather than according to the “active” or “inactive” halves of the reproductive tract (Figure S1B). In light of this, we considered these samples as replicates and merged them for downstream analysis. Wallaby gravid and non-gravid endometrium samples were also merged to make the data comparable. We analysed the ontologies of the most highly expressed genes in the monotreme reproductive tract samples (Figure S2). Overall, both species’ most significantly enriched ontologies centred around gene expression, protein synthesis, and intracellular localisation, though platypus reproductive tract gene expression was also enriched for several developmental pathways including tube morphogenesis and tissue development (Figure S2A).

**Table S1: Quality metrics for datasets used in this study.** Asterisk (*) indicates datasets generated in the present study.

| Tissue | Accession(s) | No. reads (raw) | Reads mapped (%) | Reads assigned (%) |
| --- | --- | --- | --- | --- |
| Platypus active reproductive tract* |  | 28,132,223 | 86.35 | 72.2 |
| Platypus inactive reproductive tract* |  | 27,808,946 | 85.07 | 74.4 |
| Platypus gravid endometrium | SRR1289525 | 32,805,984 | 67.62 | 76.0 |
| Platypus liver | SRR306735 | 15,860,299 | 83.26 | 79.5 |
| Platypus testis | SRR306739 | 18,898,747 | 87.46 | 71.4 |
| Echidna active reproductive tract* |  | 25,051,961 | 78.67 | 73.7 |
| Echidna inactive reproductive tract* |  | 26,723,946 | 79.96 | 75.4 |
| Echidna lactating mammary gland* |  | 25,486,293 | 84.66 | 93.9 |
|  |  | 23,503,636 | 85.47 | 89.6 |
| Echidna non-lactating mammary gland* |  | 26,522,885 | 81.41 | 80.9 |
| Echidna liver | SRR10530490 | 37,448,824 | 91.53 | 77.2 |
| Echidna testis | SRR10530485 | 26,753,678 | 93.26 | 75.8 |
| Wallaby gravid endometrium | SRR25031714 | 42,791,198 | 97.07 | 72.8 |
|  | SRR25031731 | 46,040,130 | 98.11 | 77.2 |
| Wallaby non-gravid endometrium | SRR25031721 | 47,108,516 | 96.12 | 77.8 |
|  | SRR25031720 | 46,351,944 | 97.36 | 75.1 |
| Wallaby placenta | SRR5074467 | 26,178,889 | 71.96 | 41.5 |
|  | SRR5074468 | 30,198,040 | 67.34 | 52.0 |
|  | SRR5074469 | 12,954,793 | 66.26 | 53.3 |
|  | SRR5074471 | 28,217,821 | 80.27 | 63.5 |
|  | SRR5074472 | 29,716,294 | 72.47 | 62.4 |
|  | SRR5074473 | 15,285,618 | 68.87 | 51.2 |
| Wallaby lactating mammary gland | SRR5074477 | 19,457,698 | 87.77 | 5.9 |
|  | SRR5074478 | 39,892,227 | 87.52 | 2.6 |
|  | SRR5074479 | 35,745,171 | 78.26 | 2.7 |
| Wallaby liver | SRR1041778 | 55,727,170 | 97.53 | 78.1 |
| Wallaby testis | SRR1041779 | 72,776,409 | 97.69 | 74.5 |
| Mouse gravid endometrium | SRR15417365 | 27,246,179 | 98.44 | 85.7 |
| Mouse placenta | ENCSR000BZP | - | - | 70.1 |
| Mouse lactating mammary gland | SRR1552448 | 31,425,424 | 98.33 | 81.3 |
|  | SRR1552454 | 27,225,012 | 98.35 | 85.6 |
|  | SRR1552449 | 31,276,061 | 98.46 | 81.4 |
|  | SRR1552455 | 25,433,157 | 98.20 | 86.1 |
| Mouse non-lactating mammary gland | SRR1552450 | 30,109,290 | 98.48 | 81.1 |
|  | SRR1552444 | 27,919,481 | 97.67 | 78.7 |
| Mouse liver | ENCSR216KLZ | - | - | 77.9 |
| Mouse testis | ENCSR266ESZ | - | - | 72.9 |

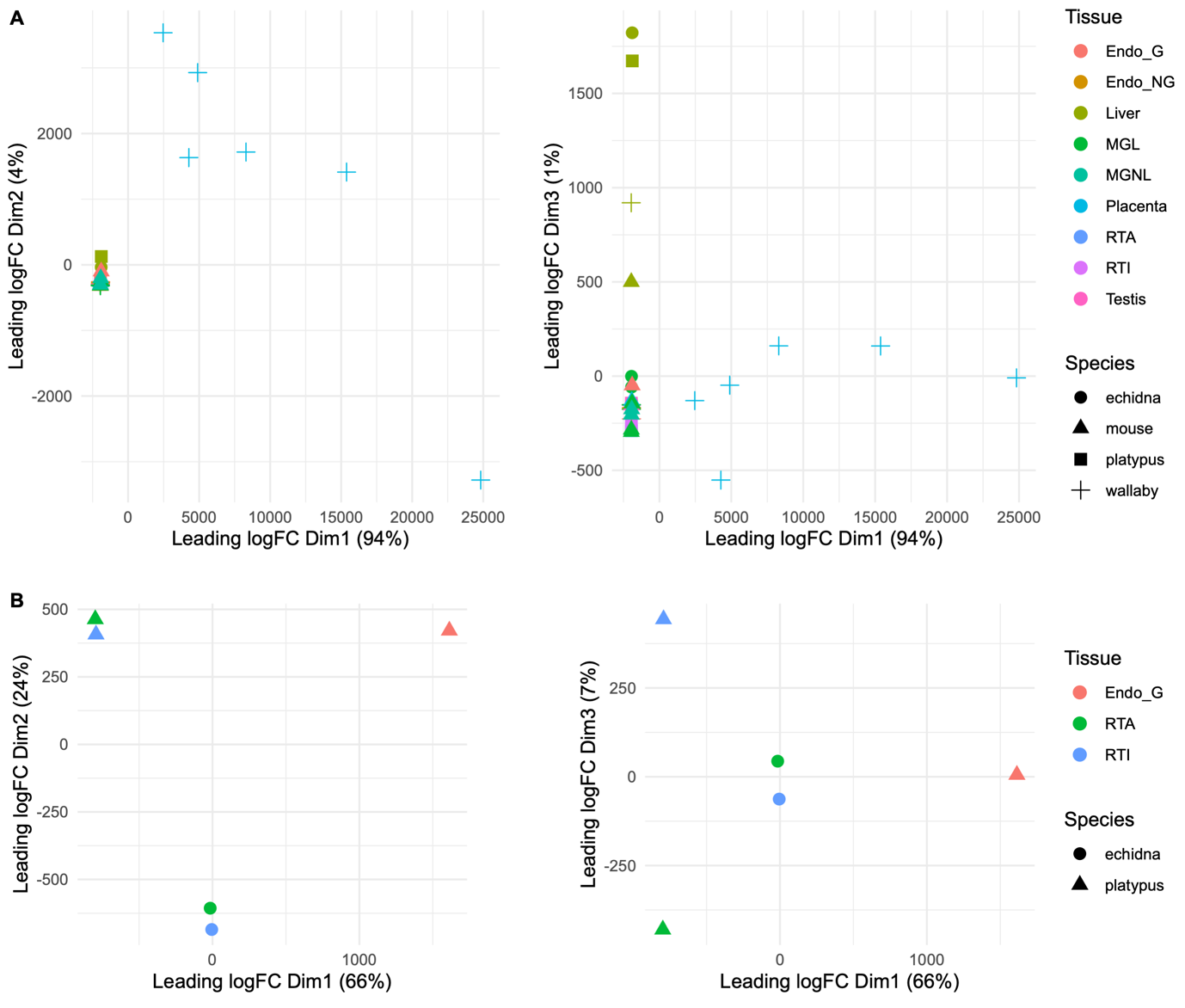

**Figure S1: MDS plotting of samples intended for analysis.** (A) All samples, showing separate clustering of wallaby placenta samples. (B) Monotreme reproductive tract datasets, showing platypus endometrium dataset clustering with echidna samples rather than platypus samples along the x-axis (dimension 1). Abbreviations: Endo_G = gravid endometrium, Endo_NG = non-gravid endometrium, MGL = lactating mammary gland, MGNL = non-lactating mammary gland, RTA = active reproductive tract, RTI = inactive reproductive tract.

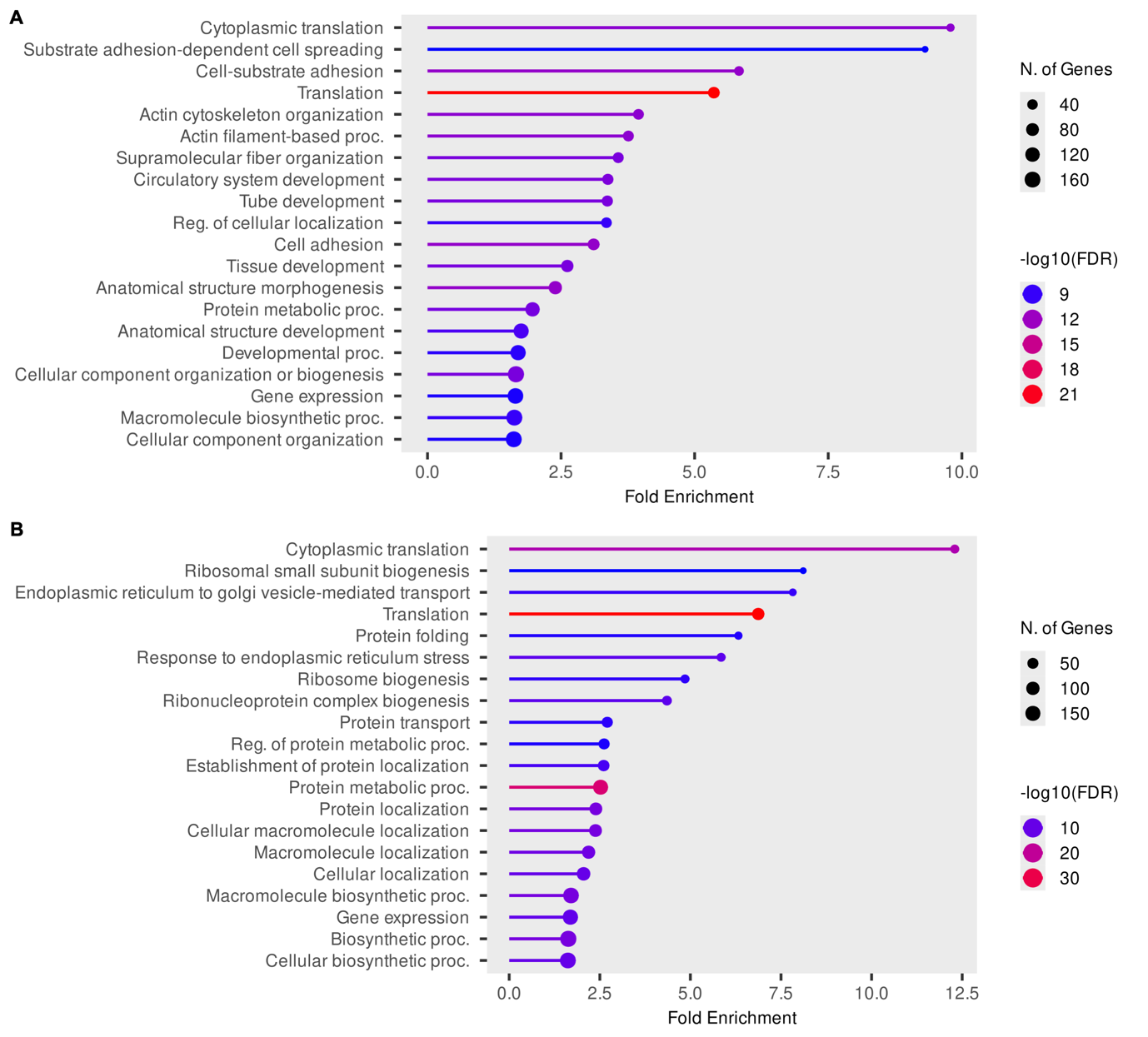

**Figure S2: Ontology of the genes most highly expressed in the (A) platypus and (B) echidna reproductive tract.** The top 500 genes (sorted by CPM) from each species were analysed using the GO Biological Process database (ShinyGO v. 0.85.1). Only the top 20 ontologies (sorted by fold enrichment) are displayed.

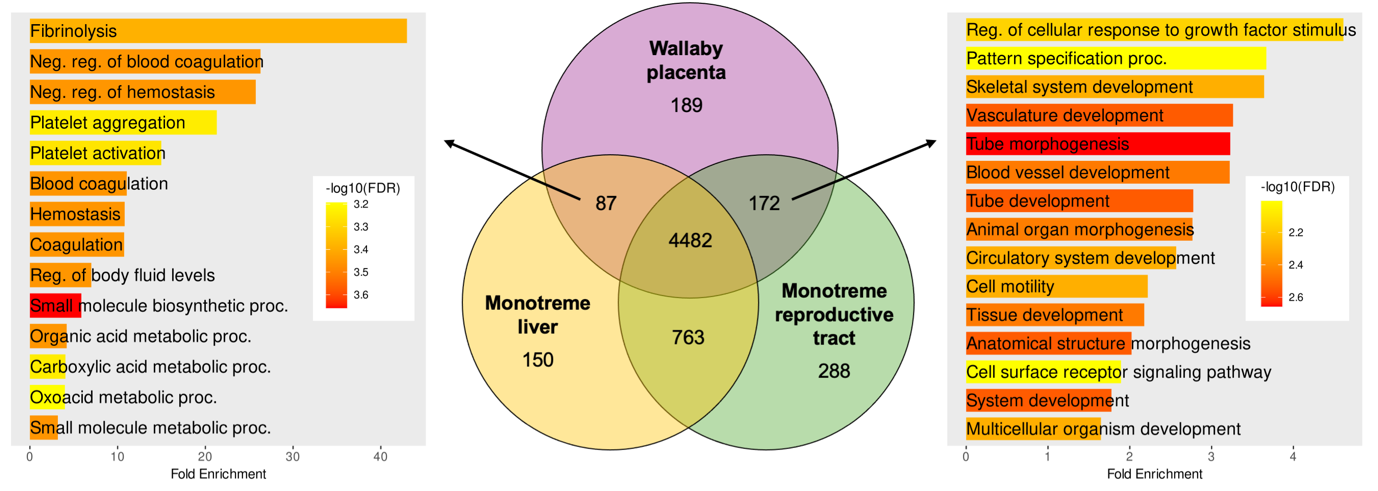
**Figure S3: Functional analysis of overlapping gene expression between the wallaby placenta, monotreme reproductive tract, and monotreme liver.** Bar plots show enrichment analysis using the GO Biological Process database; the top 15 most significant terms (ordered by fold enrichment) are displayed.

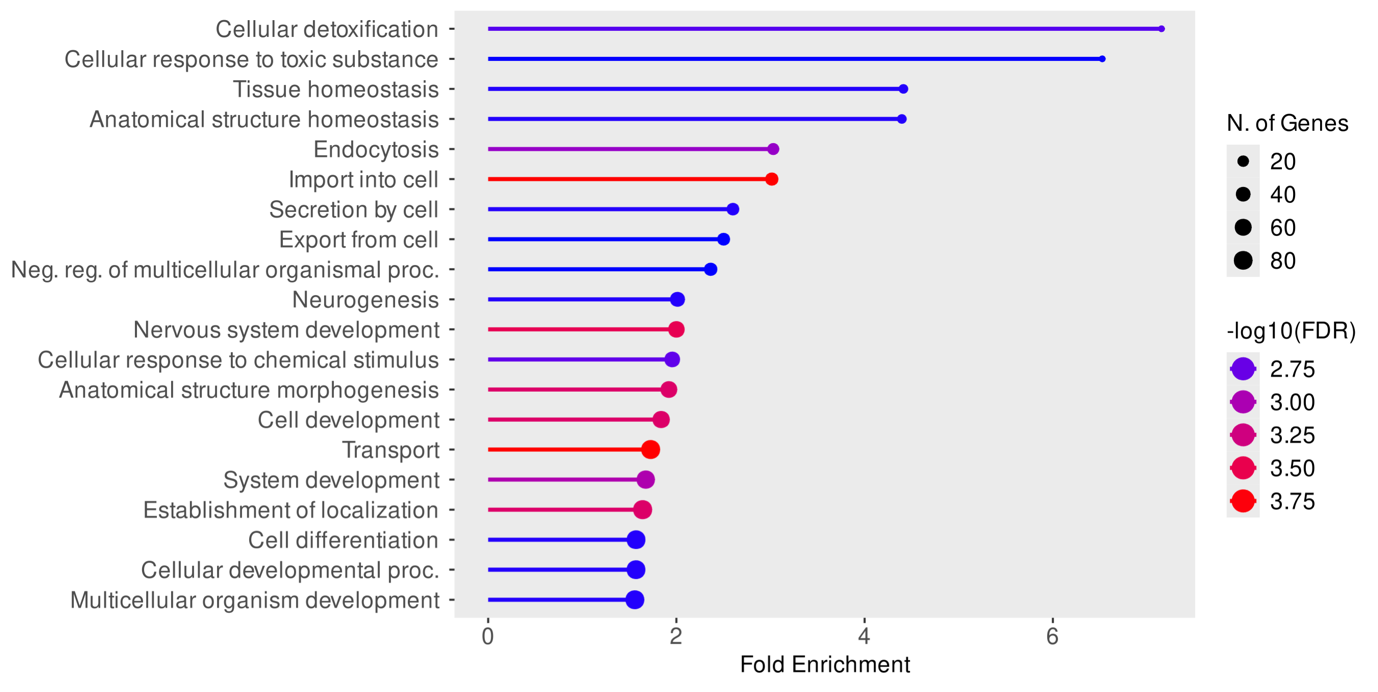

**Figure S4: Shared expression of development genes in the echidna mammary gland and mouse placenta.** Gene set enrichment analysis of genes expressed in the echidna lactating mammary gland and mouse placenta but not in the mouse lactating mammary gland. Analysis was performed using the GO Biological Process database; the top 20 most significant terms (ordered by fold enrichment) are displayed.

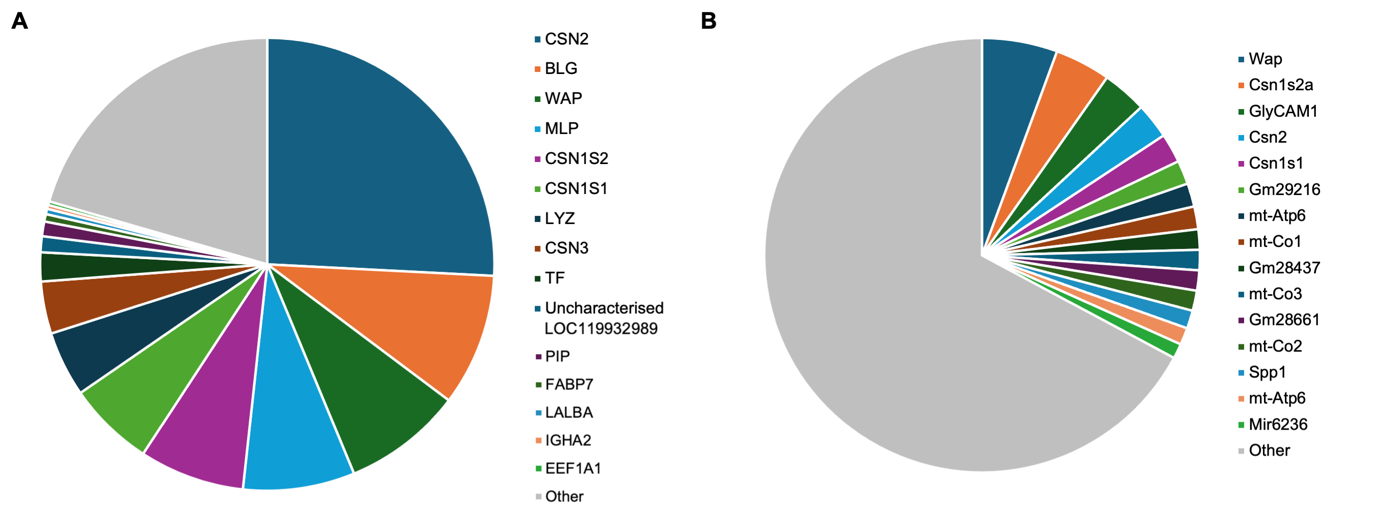
**Figure S5: Pie charts comparing gene expression in (A) echidna and (B) mouse lactating mammary gland.** Top 15 genes (sorted by TPM) are labelled individually; all remaining expressed genes are labelled “other”.

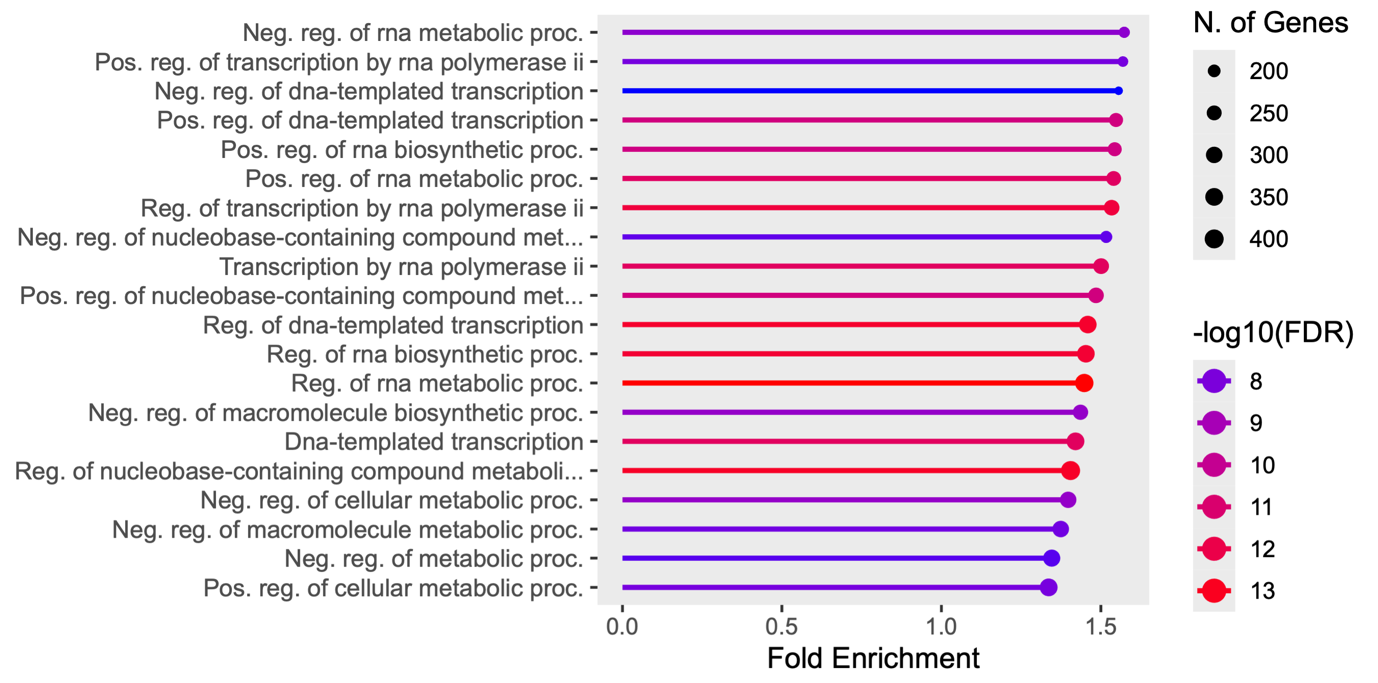

**Figure S6: Gene set enrichment analysis of mammary gland genes with lower expression in the echidna relative to the mouse using the GO Biological Process database.** Top 20 pathways are displayed in order of fold enrichment.
